# Myeloid Suclg2 deficiency attenuates aortic dissection by reshaping succinate-associated macrophage remodelling

**DOI:** 10.64898/2026.06.24.734396

**Authors:** Mingxin Xie, Shiqi Gao, Enzehua Xie, Haoyu Gao, Zhang Kai, Zhonghua Shen, Xiaogang Sun

**Affiliations:** Department of Vascular Surgery, Fuwai Hospital, National Center for Cardiovascular Diseases, Chinese Academy of Medical Sciences and Peking Union Medical College, Beijing 100037, China; Department of Cardiovascular Surgery, West China Hospital, Sichuan University, Chengdu, China; Department of Cardiology, Beijing Anzhen Hospital, Capital Medical University, National Clinical Research Center for Cardiovascular Diseases, Beijing 100029, China; Department of Cardiovascular Surgery, Second Affiliated Hospital Zhejiang University School of Medicine, Hangzhou, Zhejiang, 310001, China

**Keywords:** Aortic dissection, Suclg2, Succinate, Macrophage immunometabolism

## Abstract

**Background:** Succinate has emerged as an immunometabolic mediator of cardiovascular diseases. However, the enzymatic mechanisms linking macrophage succinate metabolism to aortic dissection remain incompletely understood. This study investigated whether Suclg2, which encodes the GDP-forming β-subunit of succinyl-CoA ligase, regulates succinate-associated macrophage remodelling and aortic dissection progression.

**Methods:** Suclg2 expression was examined in BAPN-induced AD and human acute type A aortic dissection tissues by Western Blot and immunofluorescence. Myeloid- and smooth muscle cell-specific Suclg2 conditional knockout mice were subjected to BAPN treatment to evaluate survival, aortic outcomes, histological injury and aortic morphology. Aortic RNA-seq was used to discover transcriptional changes. Bone marrow-derived macrophages were analysed under basal, M1-like and M2-like conditions to assess macrophage-intrinsic transcriptional responses. Plasma succinate levels and untargeted metabolomic profiles were further examined.

**Results:** Suclg2 was increased in murine and human dissected aortas and partially localized to CD68⁺ cells. Myeloid Suclg2 deletion markedly reduced BAPN-induced aortic rupture and dissection, whereas smooth muscle cell Suclg2 deletion did not confer comparable protection. Aortic transcriptomic analysis showed that myeloid Suclg2 deficiency attenuated inflammatory adhesion and matrix-destructive programmes. In macrophages, Suclg2 deletion did not induce a simple M1/M2 polarization shift; instead, it remodelled lipid-handling, phagolysosomal, adhesive and matrix-remodelling pathways across stimulation states. Metabolic profiling showed reduced circulating succinate and broader changes in central carbon, lipid-associated, nucleotide and redox-related metabolites after myeloid Suclg2 deletion.

**Conclusions:** Myeloid Suclg2 is a succinate-associated immunometabolic regulator of aortic dissection. Its deficiency protects against aortic dissection by reshaping macrophage inflammatory-remodelling programmes and the systemic metabolic environment.

## 1. Introduction

Acute type A aortic dissection (ATAAD) is a life-threatening vascular disease characterized by medial degeneration, inflammatory infiltration and extracellular matrix disruption^1–3^. Recent studies have highlighted metabolism as an active regulator of vascular inflammation and remodelling, rather than a passive consequence of tissue injury^4^. Succinate has emerged as a signalling metabolite with direct relevance to cardiovascular disease. Succinate signalling through SUCNR1/GPR91 has been implicated in cardiomyocyte hypertrophy, adverse cardiac remodelling and vascular inflammation^5,6^. Extracellular succinate can also amplify macrophage-associated cytokine responses and promote atherosclerotic inflammation through the SUCNR1–IL-1β axis^7^. Recently, circulating succinate was identified as both a biomarker and functional mediator of aortic aneurysm and dissection. That study showed exogenous succinate aggravated BAPN-induced AD, whereas suppression of macrophage-associated succinate production attenuated disease progression^8^.

Macrophages are highly plastic immune cells whose inflammatory and reparative functions are tightly coupled to their metabolic state. In aortic dissection, macrophages accumulate within the diseased aortic wall and contribute to inflammatory activation, protease production and extracellular matrix remodelling^9–11^. Metabolic reprogramming further shapes these macrophage responses^12,13^. Succinate can accumulate in inflammatory macrophages and act as an immunometabolite by stabilizing HIF-1α, enhancing IL-1β production and promoting mitochondrial ROS generation^14–16^.

Suclg2 encodes the GDP-forming β-subunit of succinyl-CoA ligase, a mitochondrial enzyme positioned at the succinyl-CoA–succinate step of the tricarboxylic acid cycle^17,18^. Given its metabolic location, Suclg2 may represent an underexplored node linking macrophage mitochondrial metabolism to succinate accumulation and vascular inflammatory remodelling. Whether macrophage Suclg2 regulates circulating succinate, macrophage states and AD progression remains unknown. In this study, we investigated the role of myeloid Suclg2 in BAPN-induced mouse AD and human ATAAD tissues.

## 2. Materials and Methods

### 2.1 Animals

Suclg2 flox mice were generated by Cyagen Biosciences. Mouse strains, including SMMHC-CreERT2 and Lyz2-Cre were purchased from Cyagen Biosciences. Four-week-old C57BL/6J mice were fed a custom diet containing BAPN (4 g/kg) for 4 consecutive weeks to induce Aortic Dissection. The surviving mice were anesthetized by an intraperitoneal injection of sodium pentobarbital (150 mg/kg body weight), euthanasia was confirmed by a secondary physical method (cervical dislocation) to ensure death. All experimental procedures involving animals were reviewed and approved by the Institutional Animal Care and Use Committee (IACUC) of Fuwai Hospital, Chinese Academy of Medical Sciences (Approval No. 0109-7-150-ZX(X)-42), and were conducted in strict adherence to the standards of the National Institutes of Health.

### 2.2 Cell culture

Bone marrow–derived macrophages (BMDMs) were generated from mice of the indicated genotypes (wild-type C57BL/6J and myeloid-specific Suclg2 conditional knockout mice). For bone marrow isolation, mice were deeply anesthetized with intraperitoneal pentobarbital sodium (150 mg/kg) and euthanized by cervical dislocation. Bone marrow cells were flushed from femurs and tibias and cultured in RPMI-1640 supplemented with 10% fetal bovine serum, 1% penicillin–streptomycin, and M-CSF (25 ng/mL) for 7 days to generate BMDMs. After differentiation, BMDMs were treated with LPS (100 ng/mL) to induce an M1-like state or IL-4 (20 ng/mL) to induce an M2-like state for 24 h. Culture media were refreshed every 3 days during the differentiation process

### 2.3 Statistical analysis

All statistical analyses were performed with GraphPad Prism software 9.0 (GraphPad Software). Kaplan-Meier survival curves were used to examine mouse survival rates, and the differences were analyzed with the log-rank (Mantel-Cox) test. AD incidence was analyzed with the Fisher exact test. The normality of distributions was verified by the Shapiro-Wilk test. For comparison between 2 groups, the independent t test was used for normally distributed data with equal variance, the Welch t test was used for data with unequal variances, and the Mann-Whitney test was used for nonnormally distributed data. For comparisons among ≥3 groups, 1-way ANOVA followed by the Tukey post hoc test was used for normally distributed data, and the Kruskal-Wallis test with Bonferroni post hoc tests was used when the data were nonnormally distributed. Parametric data are represented as mean±SEM, and nonparametric data are represented as median ± 95% confidence interval.

**Details of the statistical analyses are provided in the Supplemental Methods.**

## 3. Results

### 3.1 Suclg2 is elevated in AD tissues and is associated with CD68⁺ macrophage infiltration

We first examined Suclg2 expression in dissected aortic tissues. Western blot (WB) showed that Suclg2 protein was increased in aortas from BAPN-treated mice compared with WT controls (Fig. 1 A). A similar increase was observed in human ATAAD specimens (Fig. 1 B). Thus, Suclg2 upregulation was consistently detected in both mouse and human AD tissues. We next examined the spatial distribution of Suclg2 in relation to CD68⁺ cells within dissected aortic lesions. Immunofluorescence (IF) staining revealed increased CD68⁺ cell infiltration in AD aortas, accompanied by enhanced Suclg2 signal within the diseased aortic wall (Fig. 1 C-E). A subset of CD68⁺ cells showed detectable Suclg2 positivity, and quantitative analysis confirmed a higher proportion of CD68⁺Suclg2⁺ cells in AD aortas than in WT controls (Fig. 1 C). The same pattern was observed in human ATAAD tissues, where increased CD68 staining was accompanied by an elevated fraction of CD68⁺ Suclg2⁺ cells (Fig. 1 D). These data showed that Suclg2 upregulation in AD tissues is at least partly associated with infiltrating CD68⁺ macrophage/macrophage-lineage cells.

**Figure 1.**
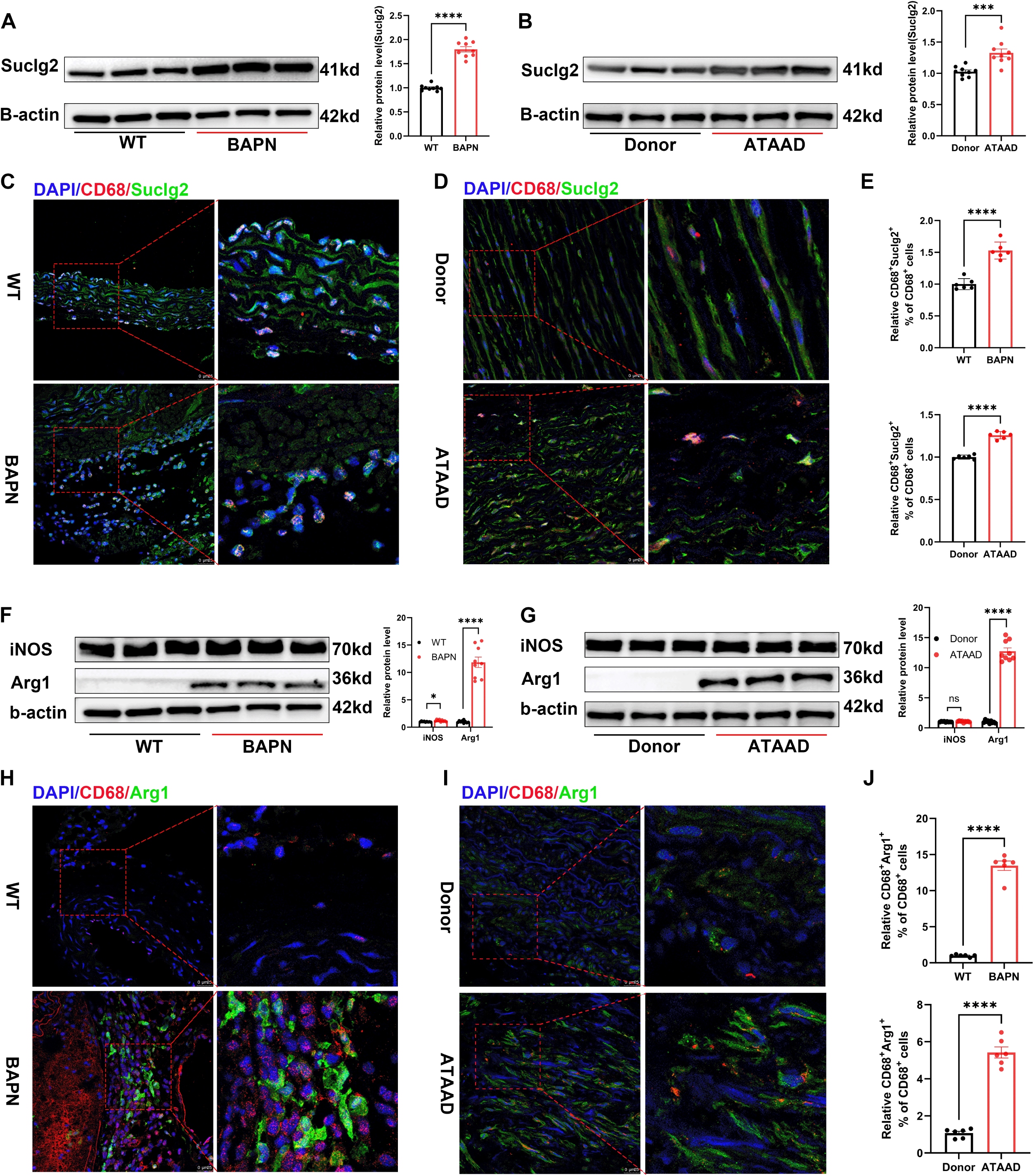
Suclg2 is increased in dissected aortas and associates with CD68⁺ macrophage. **(A, B)** Representative western blots and quantification of Suclg2 protein in mouse aortas after BAPN treatment **(A)** and in human donor (non-ATAAD) and ATAAD aortic tissues **(B)**. **(C–E)** Representative immunofluorescence staining of CD68 and Suclg2 in mouse **(C)** and human **(D)** aortic tissues, with quantification of CD68⁺ Suclg2⁺ cells among total CD68⁺ cells **(E)**. Boxed regions are shown at higher magnification. **(F, G)** Representative western blots and quantification of iNOS and Arg1 protein levels in mouse **(F)** and human **(G)** aortic tissues. **(H–J)** Representative immunofluorescence staining of CD68 and Arg1 in mouse **(H)** and human **(I)** aortic tissues, with quantification of CD68⁺Arg1⁺ cells among total CD68⁺ cells **(J)**. Boxed regions are shown at higher magnification. Data are presented as mean ± SEM. Each dot represents one biological replicate. Statistical analyses were performed as described in Methods. *P < 0.05, ***P < 0.001, ****P < 0.0001. ATAAD, acute type A aortic dissection; BAPN, β-aminopropionitrile; WT, wild type.

We further examined iNOS and Arg1, two commonly used markers associated with macrophage activation states. Across both mouse and human AD samples, Arg1 showed a more consistent increase than iNOS (Fig. 1 F, G). Co-immunofluorescence analysis further demonstrated an accumulation of CD68⁺Arg1⁺ cells in diseased aortic tissues (Fig. 1 H-J). These findings indicate that the macrophage response in ATAAD is not dominated by a uniform iNOS-high inflammatory phenotype, but is accompanied by a prominent Arg1-associated remodelling feature.

### 3.2 Myeloid Suclg2 deficiency attenuates BAPN-induced AD

To determine the contribution of Suclg2 in different cellular compartments during AD progression, Suclg2^Flox^, myeloid-specific Suclg2-deficient mice (Suclg2^Δlyz^) and vascular smooth muscle cell-specific Suclg2-deficient mice (Suclg2^ΔSMC^) were subjected to BAPN treatment. Compared with Suclg2^Flox^ mice, Suclg2^ΔLyz^ mice showed improved survival, with death numbers of 17/34 and 9/31, respectively. By contrast, Suclg2^ΔSMC^ mice did not display a comparable protective trend after BAPN treatment (13/26) (Fig. 2 A). Analysis of aortic outcomes further supported the protective effect of myeloid Suclg2 deletion. In BAPN-treated Suclg2^Flox^ mice, aortic rupture, non-ruptured TAD and lesion-free aortas accounted for 50.0%, 20.6% and 29.4% of cases, respectively. In Suclg2^ΔLyz^ mice, the rupture rate decreased to 22.6%, non-ruptured TAD decreased to 9.7%, and lesion-free aortas increased to 67.7%. In contrast, Suclg2^ΔSMC^ mice showed rupture, TAD and lesion-free aorta rates of 50.0%, 26.9% and 23.1%, respectively (Fig. 2 B). Gross morphology, H&E staining, EVG staining and ultrasound assessment were consistent with these outcome data (Fig. 2 C, D). Compared with Suclg2^Flox^ mice, Suclg2^ΔLyz^ mice exhibited less aortic dilatation, reduced medial disruption and better preservation of elastic fibre integrity. Histological scoring and ultrasound-based morphometric analysis further confirmed attenuation of BAPN-induced aortic structural damage after myeloid Suclg2 deletion (Fig. 2 E). Together, these results identify macrophage Suclg2 as a functionally relevant contributor to BAPN-induced ATAAD progression.

**Figure 2.**
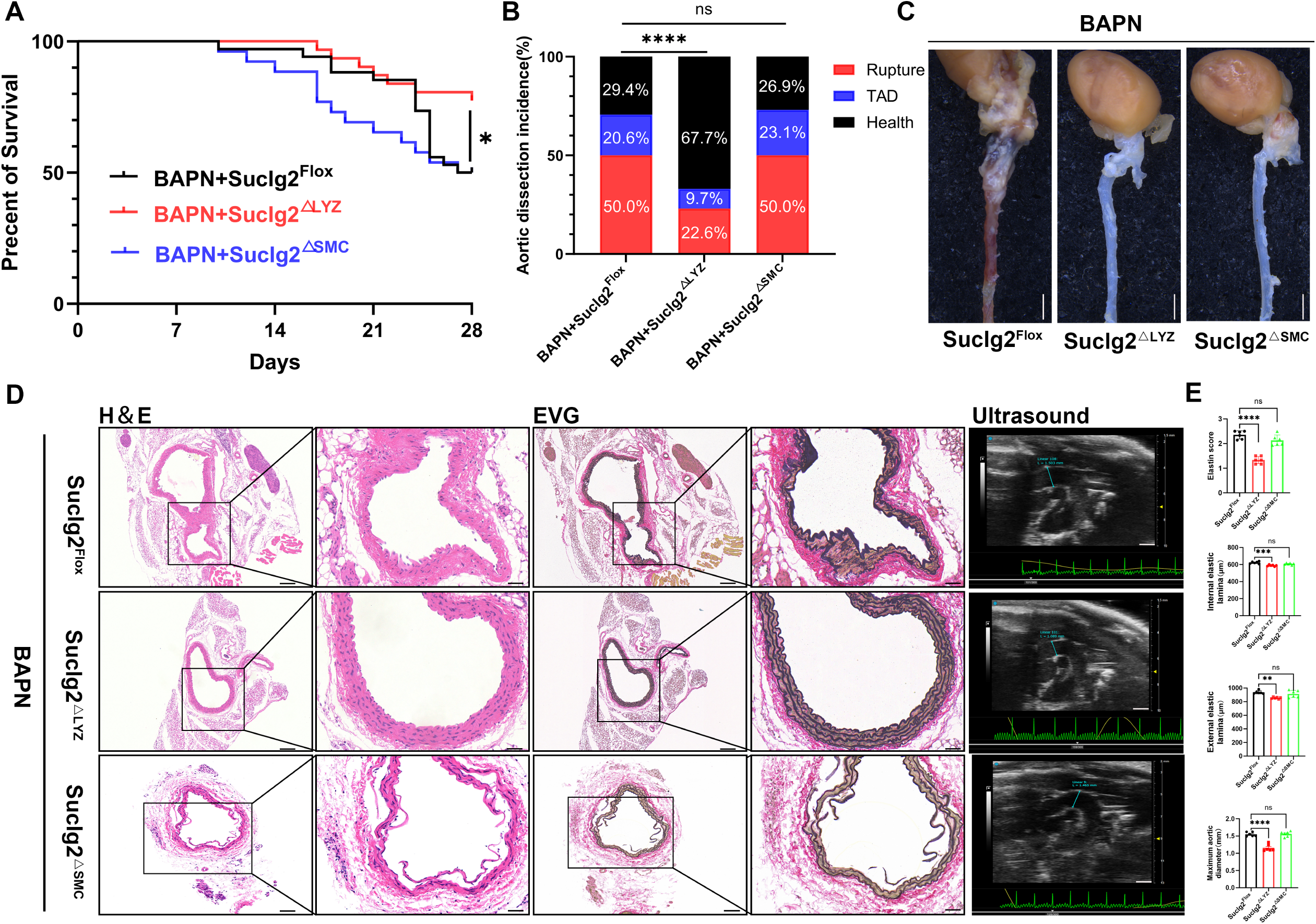
Myeloid Suclg2 deficiency attenuates aortic dissection. **(A–C)** Kaplan–Meier survival curves **(A)**, aortic outcome distribution **(B)** and representative gross images **(C)** of BAPN-treated Suclg2^Flox^, Suclg2^ΔLYZ^ and Suclg2^ΔSMC^ mice. Aortic outcomes were classified as rupture, non-ruptured TAD or lesion-free aorta. **(D, E)** Representative H&E staining, EVG staining and ultrasound images **(D)**, with quantification of elastin injury score, internal elastic lamina diameter (IEL), external elastic lamina diameter (EEL) and maximal aortic diameter **(E)** in the indicated groups. Boxed histological regions are shown at higher magnification. Data are presented as mean ± SEM. Survival was analysed by log-rank test; other statistical analyses were performed as described in Methods. *P < 0.05, **P < 0.01, ****P < 0.0001; ns, not significant. BAPN, β-aminopropionitrile; TAD, thoracic aortic dissection.

### 3.3 Myeloid Suclg2 deficiency remodels transcriptional programmes in AD aortas

To define transcriptional changes associated with myeloid Suclg2 deletion in diseased aortas, RNA-seq was performed on WT, AD and AD+Suclg2^ΔLyz^ aortic tissues (Fig. 3 E). Compared with WT aortas, AD tissues displayed canonical disease-associated transcriptional alterations, including loss of smooth muscle contractile markers, activation of extracellular matrix remodelling genes and increased inflammatory cell-associated signals. Contractile smooth muscle genes, including Acta2, Myh11, Cnn1 and Myl9, were reduced in AD aortas, whereas matrix-remodelling and matrix-degrading genes such as Col1a1, Col3a1, Thbs1, Spp1, Mmp3, Mmp13 and Plau were increased. Inflammatory recruitment-associated genes, including Tnf, Ccl3, Cxcl2 and Nlrp3, were also elevated, consistent with the coexistence of structural remodelling and inflammatory activation in the dissected aortic wall (Fig. 2 J). Direct comparison of AD+Suclg2^ΔLyz^ and AD aortas identified 90 significant differentially expressed genes, including 28 upregulated and 62 downregulated genes (Fig. 2 G). Upregulated genes included Ugt1a8, Pdk4, Msl3, Zbtb16, Stc2 and Tnfrsf25, which are linked to metabolic regulation, transport and cellular stress responses. Downregulated genes included Lyz2, Mmp3, Retnlg and Sell, representing macrophage inflammatory activation, cell adhesion and matrix-degrading programmes (Fig. 2 H). KEGG enrichment analysis showed that these differentially expressed genes were enriched in pathways related to fluid shear stress and atherosclerosis, metabolism of xenobiotics by cytochrome P450, cell adhesion molecules, IL-17 signalling, and arginine and proline metabolism (Fig. 2 I). Notably, myeloid Suclg2 deficiency did not restore smooth muscle contractile markers such as Acta2, Tagln, Myh11 and Cnn1. Instead, transcriptional changes were more prominent in inflammatory adhesion and matrix-destructive programmes. Mmp9 and Mmp13 were significantly reduced in AD+Suclg2^ΔLyz^ aortas, and several AD-induced remodelling or inflammatory genes, including Mmp3, Plaur, Lgals3, Il1b and Ccl2, showed a downward trend. Meanwhile, AD-induced phagocytic/reparative genes such as Spp1, Trem2, C1qa, Msr1 and Apoe were not uniformly abolished, but showed variable degrees of remodelling (Fig. 2 J). qPCR validation supported these RNA-seq findings. Selected smooth muscle contractile genes were not substantially restored by myeloid Suclg2 deletion, whereas representative matrix-remodelling genes, including Mmp9 and Mmp3 were reduced in AD+Suclg2^ΔLyz^ aortas compared with AD aortas (Fig. 2K). These data indicate that myeloid Suclg2 deletion primarily attenuates inflammatory adhesion and matrix-destructive transcriptional programmes while reshaping the inflammatory-remodelling balance in diseased aortas.

**Figure 3.**
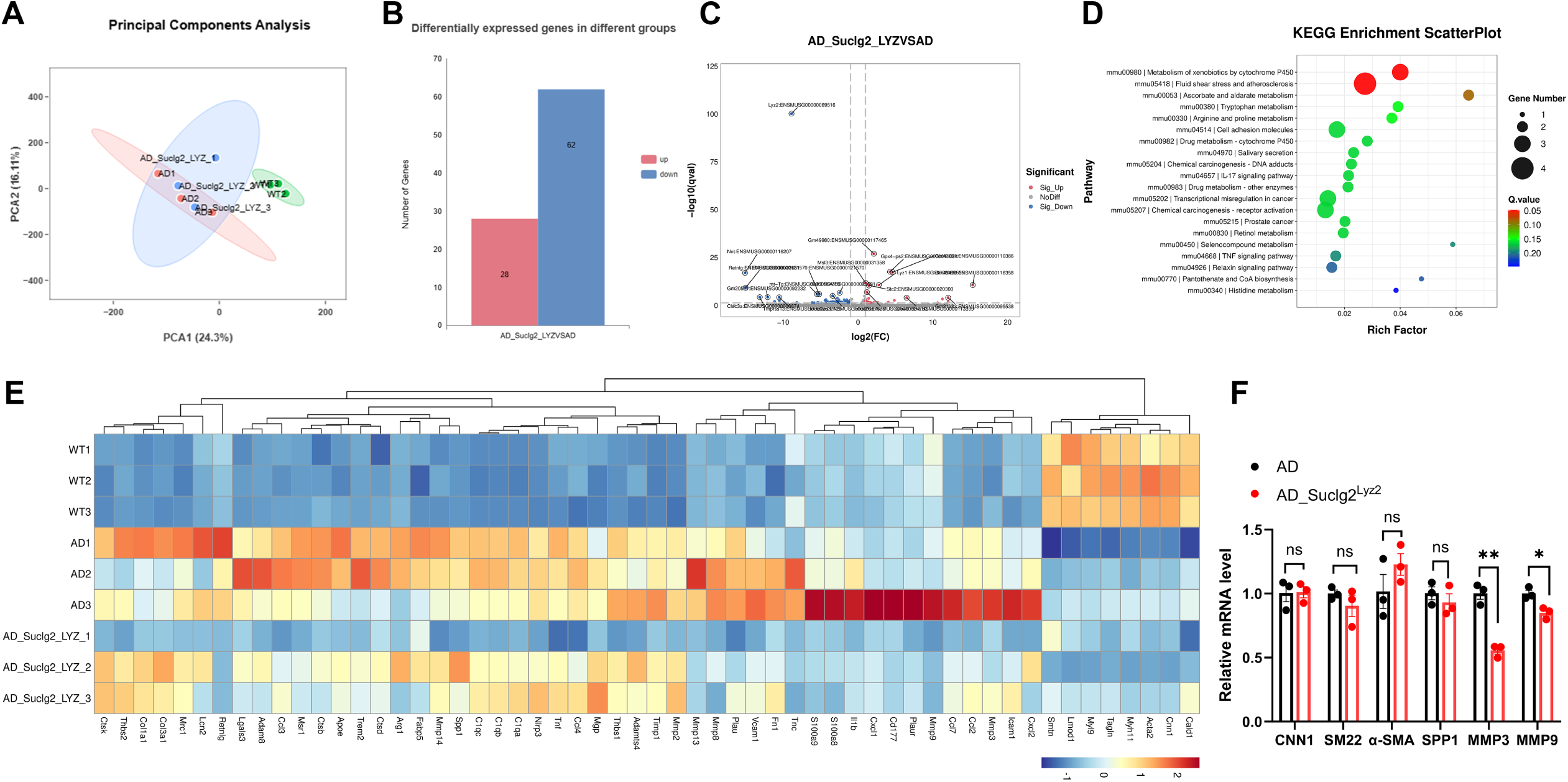
Myeloid Suclg2 deficiency reshapes transcriptional programmes in AD aortas. **(A–D)** RNA-seq analysis of aortic tissues from WT, AD and AD+Suclg2^ΔLYZ^ mice, including PCA **(A)**, the number of differentially expressed genes **(B)**, volcano plot **(C)** and KEGG pathway enrichment analysis **(D)** for AD+Suclg2^ΔLYZ^ versus AD aortas. **(E)** Heatmap of representative genes related to inflammation, adhesion, smooth muscle phenotype and extracellular matrix remodelling in WT, AD and AD+Suclg2^ΔLYZ^ aortas. **(F)** qPCR validation of contractile and remodelling-associated genes in AD and AD+Suclg2^ΔLYZ^ aortas. Heatmap values are shown as row-wise Z-scores. Data are presented as mean ± SEM. *P < 0.05, **P < 0.01. PCA, principal component analysis; AD, aortic dissection; KEGG, Kyoto Encyclopedia of Genes and Genomes.

### 3.4 Suclg2 deficiency reprogrammes BMDM transcriptional states

To determine whether Suclg2 deficiency directly alters macrophage-intrinsic transcriptional states, BMDMs from control and Suclg2-deficient mice were analysed under basal M0, LPS-induced M1-like and IL-4-induced M2-like conditions. Principal component analysis showed that BMDM samples were primarily separated according to stimulation state, while Suclg2-deficient samples remained distinguishable from their matched control samples within each condition (Fig. 4A). Differential expression analysis further showed that both polarization stimuli and Suclg2 deficiency induced broad transcriptional changes across the indicated comparisons (Fig. 4B). WB confirmed the expected induction of iNOS after LPS stimulation and Arg1 after IL-4 stimulation, indicating effective M1-like and M2-like activation in both control and Suclg2-deficient BMDMs (Fig. 4C).

**Figure 4.**
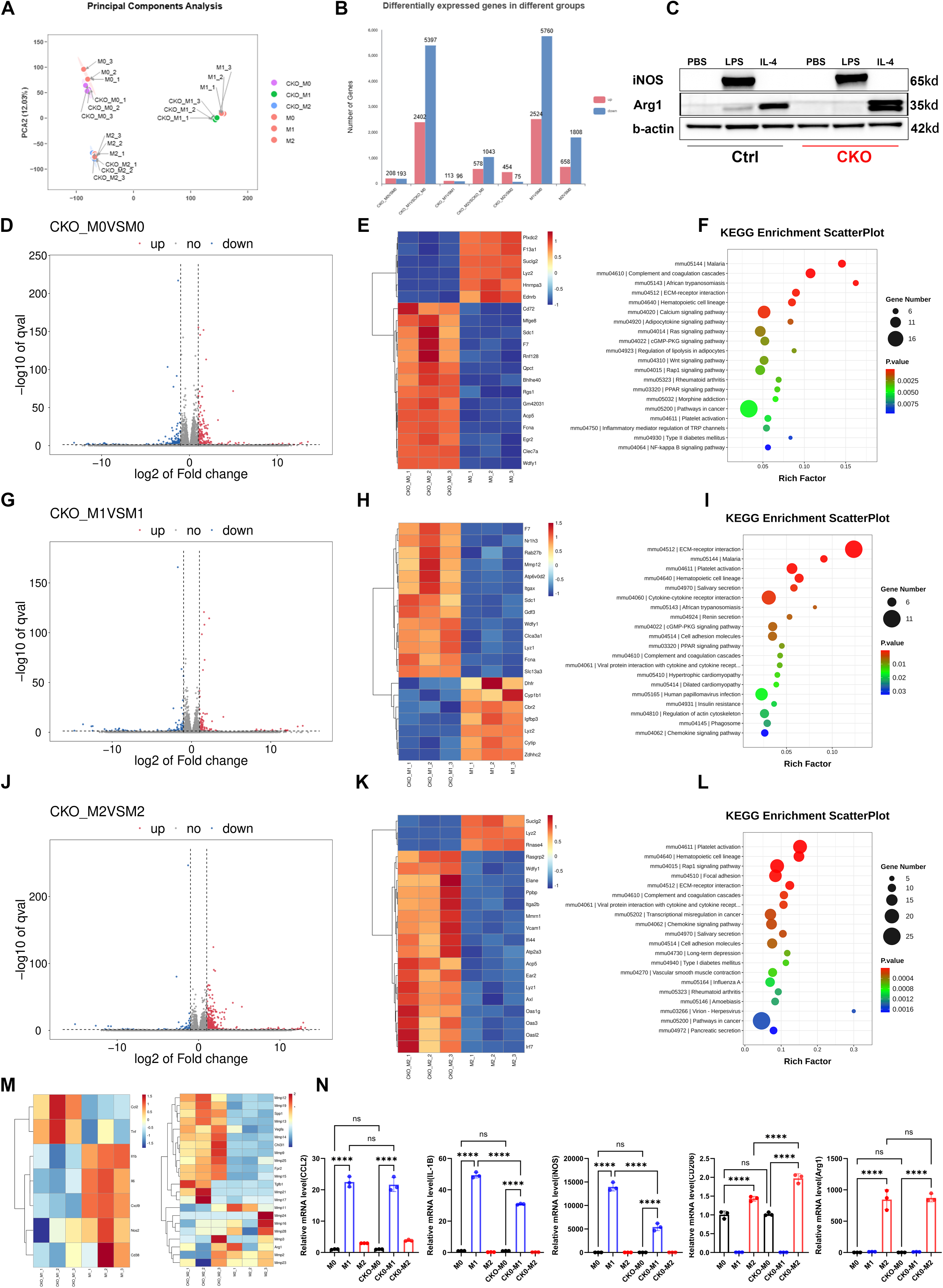
Suclg2 deficiency reprograms BMDM transcriptional states. **(A–C)** Global RNA-seq profiling and polarization validation of Flox and Suclg2-deficient BMDMs under M0, M1-like and M2-like conditions. PCA shows the distribution of RNA-seq samples **(A)**; bar plots show the number of differentially expressed genes in the indicated comparisons **(B)**; representative immunoblots show iNOS and Arg1 expression after PBS, LPS or IL-4 treatment **(C)**. **(D–L)** Volcano plots, representative heatmaps and KEGG enrichment plots are shown for each comparison of Suclg2-deficient BMDMs with matched Flox controls under M0 **(D–F)**, M1-like **(G–I)** and M2-like **(J–L)** conditions. **(M)** Heatmaps of representative inflammatory genes in M1-like BMDMs and extracellular matrix/remodelling-associated genes in M2-like BMDMs. **(N)** qPCR validation of inflammatory and reparative/remodelling-associated genes in Flox and Suclg2-deficient BMDMs under M0, M1-like and M2-like conditions. Heatmap values are shown as row-wise Z-scores. Data are presented as mean ± SEM. Statistical analyses were performed as described in Methods. *P < 0.05, ****P < 0.0001; ns, not significant. BMDM, bone marrow-derived macrophage; CKO, conditional knockout; PCA, principal component analysis; KEGG, Kyoto Encyclopedia of Genes and Genomes.

Under basal M0 conditions, Suclg2 deficiency was sufficient to alter the resting macrophage transcriptome. Volcano plot analysis identified a set of upregulated and downregulated genes in CKO-M0 BMDMs compared with control M0 BMDMs (Fig. 4D). Heatmap analysis showed increased expression of genes associated with phagolysosomal function, clearance and lipid handling, including Acp5, Atp6v0d2, Wdfy1, Clec7a, Mfge8 and lipid-associated genes such as Fabp4, Fabp5 and Plin2 (Fig. 4E). KEGG enrichment analysis showed enrichment of pathways related to extracellular matrix interaction, complement/coagulation-associated processes and metabolic or immune-related pathways (Fig. 4F). These data show that Suclg2 deficiency alters the basal macrophage state before exposure to polarizing stimuli. Under M1-like conditions, Suclg2-deficient BMDMs also displayed a distinct transcriptional profile compared with control M1-like BMDMs (Fig. 4G). Representative heatmap analysis showed increased expression of genes related to cell adhesion, phagolysosomal function, lipid handling and tissue remodelling, including Itgax, Mmp12, Atp6v0d2, Wdfy1, Sdc1 and Nr1h3 (Fig. 4H). KEGG enrichment analysis further highlighted pathways associated with cell adhesion, extracellular matrix interaction, PPAR signalling and complement/coagulation-related processes (Fig. 4I). These results indicate that Suclg2 deficiency does not abolish the response to inflammatory stimulation, but changes the transcriptional composition of M1-like macrophages.

Under M2-like conditions, Suclg2 deficiency induced the most prominent transcriptional remodelling among the matched control comparisons (Fig. 4J). The heatmap showed increased expression of innate immune and interferon-related genes, including Irf7, Ifi44, Oas1g, Mx1 and Rsad2. Genes involved in phagocytic recognition, vesicle trafficking, clearance and matrix remodelling, including Marco, Axl, Acp5, Rab27b, Wdfy1 and Mmp14, were also increased in CKO-M2 BMDMs (Fig. 4K). KEGG enrichment analysis showed enrichment of pathways related to innate immune response, response to virus, focal adhesion, ECM-receptor interaction, chemokine signalling and cell adhesion molecules (Fig. 4L). We further examined representative inflammatory and remodelling-associated gene sets. In M1-like BMDMs, the inflammatory heatmap showed a split pattern: Ccl2 and Tnf were relatively higher in Suclg2-deficient cells, whereas Il1b, Il6, Cxcl9, Nos2 and Cd38 were more prominent in control M1-like cells (Fig. 4M). In M2-like BMDMs, extracellular matrix and remodelling-associated genes showed heterogeneous changes, with several genes such as Mmp12, Mmp14, Mmp9, Spp1, Tgfb1, Vegfa and Fpr2 displaying higher or more variable expression in Suclg2-deficient cells (Fig. 4M). qPCR validation confirmed altered expression of selected inflammatory and reparative/remodelling-associated genes under M1-like and M2-like conditions (Fig. 4N).

Collectively, these results showed that Suclg2 deficiency did not drive BMDMs toward a simple M1 or M2 polarization state. Instead, Suclg2 deficiency remodeled macrophage transcriptional programmes related to phagolysosomal function, lipid handling, adhesion, innate immune sensing and extracellular matrix remodelling across basal and polarized conditions.

### 3.5 Myeloid Suclg2 deficiency alters the plasma metabolic profile in AD mice

Plasma succinic acid was first quantified in WT, AD and AD+Suclg2^ΔLyz^ mice. Compared with WT mice, AD mice showed a marked increase in plasma succinic acid peak area (171278 ± 29442 vs. 45602 ± 3898). This increase was significantly reduced in AD+Suclg2^ΔLyz^ mice (96883 ± 9648). The difference between WT and AD+Suclg2^ΔLyz^ mice was not significant (Fig. 5A). We next examined aortic expression of metabolism-related genes. The heatmap included genes involved in hypoxia signalling, glucose transport, glycolysis, lactate metabolism and TCA cycle-associated metabolism. AD and AD+Suclg2^ΔLyz^ aortas displayed different expression patterns across these genes, indicating that myeloid Suclg2 deletion was accompanied by altered metabolic gene expression in diseased aortic tissues (Fig. 5 B).

**Figure 5.**
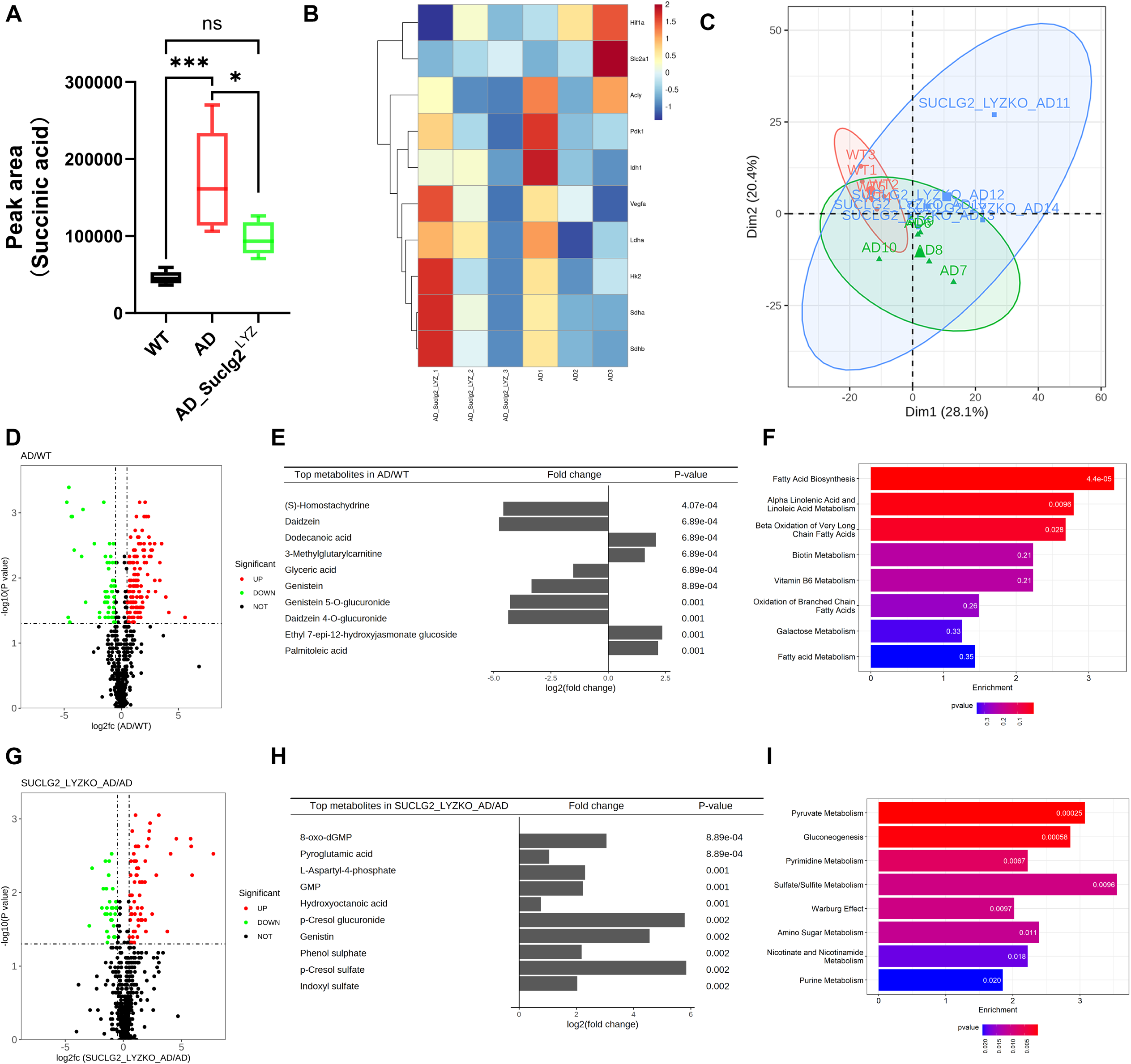
Suclg2 deficiency lowers circulating succinate and reshapes systemic metabolic profiles in AD. **(A)** Plasma succinic acid abundance in WT, AD and AD+Suclg2^ΔLYZ^ mice (n=5). **(B)** Heatmap of representative immunometabolic genes in AD and AD+Suclg2^ΔLYZ^ aortas. **(C)** PCA of plasma metabolomic profiles from WT, AD and AD+Suclg2^ΔLYZ^ mice. **(D–F)** Differential metabolomic analysis of AD versus WT plasma samples, including volcano plot **(D)**, top-ranked differential metabolites with fold change and P value **(E)**, and pathway enrichment analysis **(F)**. **(G–I)** Differential metabolomic analysis of AD+Suclg2^ΔLYZ^ versus AD plasma samples, including volcano plot **(G)**, top-ranked differential metabolites with fold change and P value **(H)**, and pathway enrichment analysis **(I)**. Heatmap values are shown as row-wise Z-scores. Data are presented as mean ± SEM. Statistical analyses were performed as described in Methods. *P < 0.05, ***P < 0.001; ns, not significant. PCA, principal component analysis; WT, wild type; AD, aortic dissection; TCA, tricarboxylic acid.

Untargeted plasma metabolomics was then performed in WT, AD and AD+Suclg2^ΔLyz^ mice. PCA showed that WT, AD and AD+Suclg2^ΔLyz^ samples displayed partially separated but overlapping distributions. The first two principal components explained 28.1% and 20.4% of the total variance, respectively (Fig. 5C). Differential metabolite analysis between AD and WT mice identified a broad set of altered metabolites (Fig. 5D). Among the top-ranked metabolites in AD versus WT, dodecanoic acid, 3-methylglutarylcarnitine, ethyl 7-epi-12-hydroxyjasmonate glucoside and palmitoleic acid were increased, whereas (S)-homostachydrine, daidzein, glyceric acid, genistein, genistein 5-O-glucuronide and daidzein 4-O-glucuronide were decreased (Fig. 5E). Pathway enrichment analysis of AD-associated differential metabolites identified fatty acid biosynthesis, α-linolenic acid and linoleic acid metabolism among the top-ranked pathways (Fig. 5F). We further compared AD+Suclg2^ΔLyz^ with AD mice to define metabolic changes associated with myeloid Suclg2 deletion in the AD setting. Volcano plot analysis showed both increased and decreased metabolites in AD+Suclg2^ΔLyz^ mice relative to AD mice (Fig. 5G). The top-ranked altered metabolites included 8-oxo-dGMP, pyroglutamic acid, L-aspartyl-4-phosphate, GMP, hydroxyoctanoic acid, p-cresol glucuronide, genistin, phenol sulphate, p-cresol sulfate and indoxyl sulfate (Fig. 5H). Pathway enrichment analysis showed that these differential metabolites were mainly associated with pyruvate metabolism, gluconeogenesis, pyrimidine metabolism and sulfate/sulfite metabolism (Fig. 5I). Together, these results show that AD is associated with increased circulating succinate and broad plasma metabolic alterations, while myeloid Suclg2 deletion reduces plasma succinate and is accompanied by distinct changes in central carbon, nucleotide, sulfate-related and lipid-associated metabolic pathways.

## 4. Discussion

This study identifies myeloid Suclg2 as a metabolic regulator of aortic dissection. The main advance of this study is the identification of myeloid Suclg2 as a functional metabolic node in AD, rather than the observation of succinate elevation alone, which has been reported previously. Together with reduced circulating succinate and macrophage transcriptional rewiring, these findings suggest that Suclg2 links macrophage mitochondrial metabolism to the inflammatory-remodelling environment of the dissected aortic wall.

Previous work has established succinate as an immunometabolic signal with broad relevance to cardiovascular pathology^4,19^. In heart failure, elevated circulating succinate has been linked to ATP5A1 succinylation, mitochondrial dysfunction, impaired energy metabolism and cardiomyocyte injury^20^. Beyond cardiomyocytes, extracellular succinate can signal through SUCNR1/GPR91 to modulate vascular inflammation, macrophage cytokine responses and angiogenesis-related endothelial signalling^5^. In macrophages, succinate is not only a TCA cycle intermediate but also a functional regulator of activation state, acting through HIF-1α stabilization, IL-1β induction and mitochondrial ROS production^21–23^. Recently, a study demonstrated circulating succinate was increased in aortic aneurysm and dissection, and exogenous succinate aggravated AD in mice, which linked macrophage p38α–CREB–OGDH signalling to inflammatory factor expression and succinate production^8^. Our findings identifying macrophage Suclg2 as a succinate-associated enzymatic node in AD. The reduction in plasma succinate after macrophage Suclg2 deletion, together with the protection from BAPN-induced rupture and dissection, suggests that macrophage-lineage succinate metabolism contributes to the metabolic and inflammatory-remodelling environment of the diseased aorta.

Although succinate has been studied in AD, the role of Suclg2 in aortic disease has been largely unexplored. Suclg2 encodes the GDP-forming β-subunit of succinyl-CoA ligase. Suclg2 relevant not only to succinate abundance, but also to succinyl-CoA utilization and potentially succinylation-related metabolic regulation^24,25^. Previous studies mainly focused on succinate accumulation, SDH-dependent succinate oxidation and succinylation, whereas the contribution of Suclg2 has received much less attention^26,27^. In AD, prior metabolomic and mechanistic work has established succinate as a disease-relevant metabolite and has linked macrophage-associated succinate production to disease progression^8^. Our study extends this framework by identifying macrophage Suclg2 as a succinate-associated enzymatic node associated with circulating succinate elevation and macrophage inflammatory-remodelling states during AD. Macrophage Suclg2 deletion reduced plasma succinate and protected against BAPN-induced AD suggests that Suclg2-dependent metabolism in macrophage cells contributes to the succinate-enriched disease environment. By contrast, smooth muscle cell Suclg2 deletion did not produce a comparable protective effect in our model. This indicates that the disease-modifying effect of Suclg2 is more closely linked to the macrophage immunometabolic context in BAPN-induced AD.

Macrophage Suclg2 deletion did not markedly restore the smooth muscle contractile programme in diseased aortas, suggesting that its protective effect was unlikely to result from direct recovery of smooth muscle contractile phenotype. Instead, the main transcriptional changes involved inflammatory adhesion and matrix-remodelling pathways. This is consistent with the known role of macrophages in shaping the proteolytic and reparative environment of the dissected aortic wall: macrophage-derived cytokines, chemokines and MMPs can promote leukocyte recruitment, elastic fibre fragmentation and matrix degradation, whereas distinct macrophage states may also participate in debris clearance, fibrotic repair and tissue stabilization^28–30^. Single-cell studies of ATAAD further support the presence of heterogeneous macrophage populations rather than a simple M1/M2 state, identifying inflammatory macrophages enriched for cytokine and chemokine signalling, lipid-associated expressing Apoe and phagocytic programmes, and matrix-remodelling populations characterized by extracellular matrix and protease-related genes^31,32^. In our data, macrophage Suclg2 deletion reduced matrix-destructive genes such as Mmp9 and Mmp13, while phagocytic or reparative markers including Spp1, Trem2, C1qa, Msr1 and Apoe were not uniformly abolished. Thus, Suclg2 deletion appears to rebalance macrophage-associated inflammatory-remodelling programmes, reducing proteolytic and inflammatory pressure without globally suppressing macrophage activity.

The BMDM transcriptomic data provide macrophage-intrinsic evidence that Suclg2 deficiency does not act through a simple M1/M2 polarization switch. Although the increased Arg1⁺CD68⁺ expression in AD lesions indicates an Arg1-associated macrophage response, Arg1 is increasingly recognized as a marker that can be induced in multiple macrophage states in vivo besides canonical M2 phenotype^33^. This is supported by our in vitro data. In BMDMs, Suclg2 deletion increased genes related to phagolysosomal function, lipid handling and clearance. Under LPS stimulation, the inflammatory response was not globally blocked; instead, some of chemokine and TNF-associated signals were preserved, whereas Il1b, Nos2/Cd38 and IFN-inducible chemokine modules were relatively reduced. After IL-4 treatment, Suclg2 deletion did not simply amplify a classical Arg1/Mrc1/Retnla programme, but enhanced genes related to innate immune sensing, vesicle trafficking, clearance and matrix remodelling. These findings are consistent with the current view that macrophage states in diseased tissues are heterogeneous and shaped by local metabolic and injury-related cues^34–36^. Thus, Suclg2 deficiency appears to remodel macrophage lipid-handling, phagolysosomal and matrix-remodelling programmes, rather than driving macrophages along a linear M1-or-M2 states.

The metabolomic data place macrophage Suclg2 within the systemic metabolic remodelling associated with AD. Targeted metabolite analysis showed reduced plasma succinate in AD+Suclg2^ΔLyz^ mice, consistent with the function of Suclg2 at the succinyl-CoA–succinate node of mitochondrial metabolism. However, the reduction in succinate should be interpreted as one component of a broader Suclg2-dependent metabolic reconfiguration, rather than as an isolated determinant of protection. Suclg2 functions within a reversible mitochondrial reaction and may influence succinyl-CoA/succinate flux, mitochondrial respiration, redox balance, amino acid metabolism and succinylation-related regulation^17,37^. Consistent with this broader metabolic role, untargeted plasma metabolomics revealed coordinated changes in central carbon metabolism, lipid-related pathways and redox-associated metabolites. These changes may arise from macrophage-intrinsic metabolic rewiring, from reduced disease burden, or from both processes. The present data also do not resolve the relative contribution of intracellular succinate metabolism and signalling pathway to the protective phenotype. Future studies using isotope tracing, macrophage-resolved metabolomics, succinate rescue will be required to define how Suclg2-dependent metabolic flux contributes to AD progression.

Several limitations should be acknowledged. First, although Suclg2 was increased in diseased aortic tissues and partially colocalized with CD68⁺ cells, whole-aorta immunoblotting cannot define the dominant cellular source of Suclg2 induction. Other aortic cell types, including vascular smooth muscle cells, endothelial cells and fibroblasts, may also contribute to tissue-level Suclg2 expression. Second, although Suclg2-deficient BMDMs showed robust transcriptional rewiring, in vitro macrophage stimulation cannot fully reproduce the mechanical stress, hypoxic milieu, matrix remodelling and multicellular interactions present in dissected aortic lesions. Another limitation is that the metabolic mechanism remains incompletely resolved. The reduction in circulating succinate after macrophage Suclg2 deletion supports a link between Suclg2-dependent macrophage metabolism and the AD-associated metabolic state. However, because plasma metabolomics provides a systemic readout, it cannot by itself determine whether these metabolic changes arise directly from altered macrophage metabolism or secondarily from reduced aortic injury and inflammation. Moreover, the present data do not define whether protection is mediated mainly by altered succinyl-CoA/succinate flux, intracellular succinate-dependent inflammatory signalling, mitochondrial redox changes, succinylation-related regulation or extracellular succinate signalling. Future studies will be required to define the specific causal metabolic mechanisms by macrophage Suclg2 regulates AD progression.

## 5. Conclusion

In summary, our study identifies macrophage Suclg2 as a regulator of succinate-associated immunometabolism in AD. Its protective effect is not explained by a simple M1/M2 switch or by restoration of smooth muscle contractile phenotype. Instead, macrophage Suclg2 deletion reduces circulating succinate and reshapes macrophage lipid-handling, phagolysosomal, adhesive and matrix-remodelling programmes. These changes are associated with reduced inflammatory and matrix-destructive pressure in the diseased aortic wall. Our findings provide a metabolic perspective on macrophage-driven aortic injury and suggest that succinate-associated macrophage pathways may represent potential therapeutic targets in ATAAD.

## 6. Acknowledgements

None.

## 7. Funding sources

This work was supported by the National High-Level Hospital Construction Project of Fuwai Hospital, Chinese Academy of Medical Sciences (No. 2025-GSP-GG-15).

## 8. Statement of author contributions

Mingxin Xie and Shiqi Gao participated in the Conceptualization, Data curation, Formal analysis, Methodology, Validation, Visualization, Writing – original draft, and Writing – review and editing. Enzehua Xie and Haoyu Gao participated in the Investigation, Validation and Formal analysis. Zhang Kai, Zhonghua Shen and Xiaogang Sun participated in the Investigation, Funding acquisition, Supervision, and Writing – review and editing.

## 9. Conflict of Interest Statement

The authors report no conflict of interest relative to this work.

## 10. Data availability

The data that support the findings of this study are available from the corresponding author upon reasonable request.

## Notes

### Competing Interest Statement

The authors have declared no competing interest.

